# Cell wall remodeling drives engulfment during *Bacillus subtilis* sporulation

**DOI:** 10.1101/087858

**Authors:** Nikola Ojkic, Javier López-Garrido, Kit Pogliano, Robert G. Endres

## Abstract

When starved, the Gram-positive bacterium *Bacillus subtilis* forms durable spores for survival. Sporulation initiates with an asymmetric cell division, creating a large mother cell and a small forespore. Subsequently, the mother cell membrane engulfs the forespore in a phagocytosis-like process. However, the force generation mechanism for forward membrane movement remains unknown. Here, we show that membrane migration is driven by cell wall remodeling at the leading edge of the engulfing membrane, with peptidoglycan synthesis and degradation mediated by penicillin binding proteins in the forespore and a cell wall degradation protein complex in the mother cell. We propose a simple model for engulfment in which the junction between the septum and the lateral cell wall moves around the forespore by a mechanism resembling the ‘template model’. Hence, we establish a biophysical mechanism for the creation of a force for engulfment based on the coordination between cell wall synthesis and degradation.

## Introduction

To survive starvation, the Gram-positive bacterium *Bacillus subtilis* forms durable endospores [1]. The initial step of sporulation is the formation of an asymmetrically positioned septum (polar septation), which produces a larger mother cell and a smaller forespore (Figure 1A). After division, the mother cell engulfs the forespore in a phagocytosis-like manner. Engulfment entails a dramatic reorganization of the sporangium, from two cells that lie side by side to a forespore contained within the cytoplasm of the mother cell. The internalized forespore matures and is ultimately released to the environment upon mother cell lysis. After engulfment, the forespore is surrounded by two membranes within the mother cell cytoplasm, sandwiching a thin layer of peptidoglycan (PG) [2]. While a number of molecular players for engulfment have been identified, the mechanism of force generation to push or pull the mother cell membrane around the forespore remains unknown [3].

**Figure 1:**
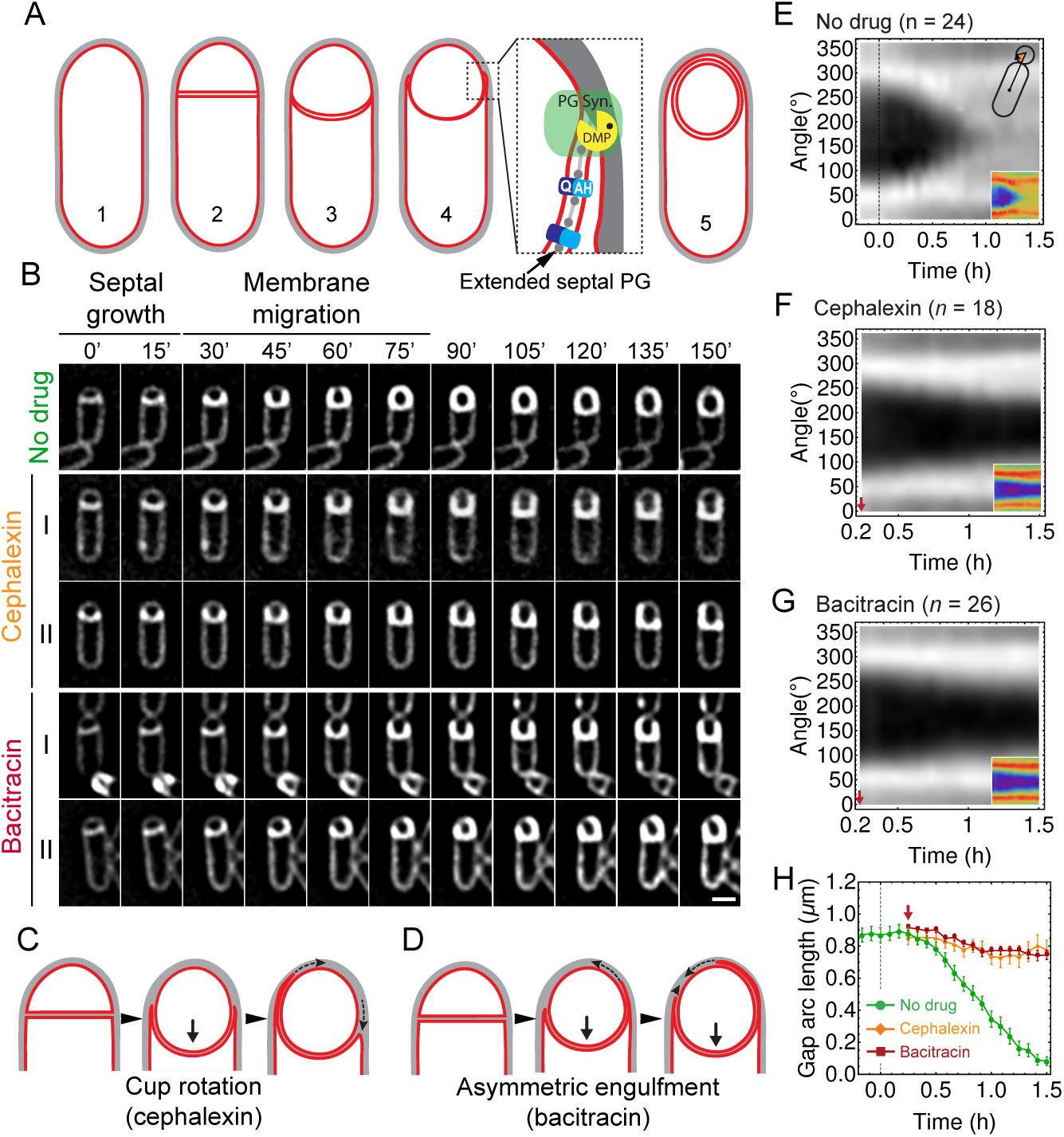
Peptidoglycan (PG) synthesis is essential for leading-edge (LE) migration. (A) Morphological changes during spore formation. Peptidoglycan shown in grey, membrane in red. (1) Vegetative cell. (2) The first morphological step in sporulation is asymmetric cell division, producing a smaller forespore and a larger mother cell. (3) The septum curves and protrudes towards the mother cell. (4) The mother cell membrane migrates towards the forespore pole. The different modules contributing to membrane migration are shown in the inset (see Introduction for details). During engulfment, the septal PG is extended around the forespore [2]. (5) Fully engulfed forespore surrounded by two membranes sandwiching a thin layer of PG. (B) Snapshots of engulfing sporangia from time-lapse movies in the absence of antibiotics, or in the presence of cephalexin or bacitracin. Cells were stained with fluorescent membrane dye FM 4-64 and imaged in medial focal plane. In the absence of antibiotics (top) the septum curves and grows towards the mother cell without significant forward movement of the engulfing membrane for ~20 min. After that, the LE of the engulfing membrane starts migrating and reaches the forespore pole in ~1 h. When PG precursor delivery system is blocked with bacitracin (50 *μg*/ml): (I) LE migration is stopped or (II) engulfment proceeds asymmetrically. Similar results are obtained when cells are treated with cephalexin (50 μg/ml). However, in this case the asymmetric engulfment phenotype observed at later time points is due to rotation of the engulfment cup (C) rather than to asymmetric movement forward of the engul ng membrane (D). (E) FM 4-64 average kymograph of *n* = 24 engulfing cells. Average fluorescent intensity along forespore contour vs time in the mother-forespore reference frame as shown in top inset. All cells are aligned in time based on time 0’ (0 min). Time 0‛ is assigned to the onset of curving septum (Figure 1-figure supplement 3). Bottom inset is average kymograph represented as heat map. (F-G) Average kymograph for cells treated with cephalexin (*n* = 18) (F) or bacitracin (*n* = 26)(G). When drug was added analyzed cells had (55 ± 5) % engulfment (red arrow). The percentage of engulfment is calculated as total angle of forespore covered with mother membrane divided by full angle. All cells had fully curved septum. Non-engulfed part of the forespore is represented as the black regions in kymographs. (H) In untreated sporangia, gap starts to close ~20 min after onset of membrane curving. In antibiotic-treated cells gap does not close. Sample size as in (F-G). Red arrow points when drug is added. Average ± SEM. Scale bar 1 *μ*m.

The cellular machinery for engulfment is complex, presumably to add robustness for survival (Figure 1A, inset). First, the forespore protein SpoIIQ and the mother cell protein SpoIIIAH interact in a zipper-like manner across the septum [4], and mediate the fast engulfment observed in the absence of cell wall [5, 6]. This complex is static and is proposed to act as a Brownian ratchet to prevent backwards movement of the engulfing membrane, contributing to the robustness of engulfment in intact cells [5, 7]. Second, the SpoIID, SpoIIM and SpoIIP complex (DMP) localizes at the leading edge (LE) of the mother cell engulfing membrane and is essential and rate limiting for membrane migration [8, 9]. The complex contains two enzymes that degrade PG in a processive manner: SpoIIP removes stem peptides, and SpoIID degrades the resulting denuded glycan strands [8–11]. Mutants with reduced SpoIID or SpoIIP activity or protein levels engulf asymmetrically, with the engulfing membrane migrating faster on one side of the forespore [8, 9]. Third, blocking PG precursor synthesis with antibiotics impairs membrane migration in mutants lacking the Q-AH zipper, suggesting that PG synthesis at the LE of the engulfing membrane contributes to engulfment [2, 12]. However, the mechanistic details of membrane migration and for the coordination between PG synthesis and degradation remain unclear.

The biophysical principles of cell wall remodeling in Gram-positive bacteria are not well understood. In *Bacillus subtilis*, the cell wall is about 20-40 nm thick, and is likely organized into multiple (20-30) PG layers [11, 13–16]. In contrast, cryo-electron tomography has demonstrated that a thin PG layer is present between the septal membranes throughout engulfment, appearing to form continuous attachments with the old cell wall [2, 17]. The outer cell wall of Gram-positive bacteria also contains a significant amount of teichoic acids, important for cell morphology, phosphates, and antibiotic resistance [18, 19] but largely absent in spores [20, 21]. Engulfment entails extensive cell wall remodeling, with peptidoglycan precursors, newly synthesized PG and the sporulation specific PG degradation machinery localizing at the LE of the engulfing membrane [2, 8, 12]. However, since engulfment occurs at high turgor pressure within the cramped confines of the thick outer cell wall, we expect that membrane movement is severely reduced by steric hindrance [22]. Hence, we anticipate that peptidoglycan remodeling is a critical step in engulfment, which may either act as a force generator or simply create room for engulfment by the mother cell membrane.

Here, we provide a biophysical mechanism for engulfment in which PG synthesis and degradation move the junction between the septal PG and the lateral cell wall around the forespore, making room for the engulfing membrane to move by entropic forces. Using antibiotics that block different steps in PG synthesis, we demonstrate that PG synthesis is essential for membrane migration in all conditions and contributes to the localization of SpoIIDMP at the LE. We also show that components of the PG biosynthetic machinery, including several penicillin binding proteins (PBPs) and the actin-like proteins MreB, Mbl and MreBH track the LE of the engulfing membrane when produced in the forespore, but not when produced in the mother cell. We implement a biophysical model for PG remodeling at the LE of the engulfing membrane, based on the ‘template mechanism’ of vegetative cell growth and implemented by stochastic Langevin simulations. These simulations reproduce experimentally observed engulfment dynamics, forespore morphological changes, and asymmetric engulfment when PG synthesis or degradation is perturbed. Taken together, our results suggest that engulfment entails coordination of PG synthesis and degradation between the two compartments of the sporangium, with forespore-associated PBPs synthesizing PG ahead of the LE and the mother-cell DMP complex degrading this PG to drive membrane migration.

## Results

### PG synthesis is essential for membrane migration

In contrast to previous studies [12], we attempted to find conditions that completely blocked PG synthesis in sporulating cultures (Figure 1-figure supplement 1). To estimate the sporulation minimal inhibitory concentration (sMIC) of antibiotics, we monitored the percentage of cells that had undergone polar septation over time in batch cultures. Polar septation depends on PG synthesis and is easy to track visually [23], which makes it a good indicator for efficient inhibition. We assayed nine antibiotics inhibiting different steps in the PG biosynthesis pathway, and found concentrations that blocked the formation of new polar septa for seven of them (Figure 1-figure supplement 1B, C). In most cases, the antibiotic concentration that blocked polar septation also inhibited completion of engulfment (Figure 1-figure supplement 1B). Only two drugs, fosfomycin and D-cycloserine, failed to completely block polar cell division. These drugs inhibit production of PG precursors that, during starvation conditions, might be obtained by recycling rather than *de novo* synthesis [13], potentially from cells that lyse during sporulation, as has been observed in studies of *B. subtilis* cannibalism [24–26], or from cells that lyse due to antibiotic treatment [27]. These results demonstrate that the later stages in PG synthesis are essential for engulfment in wild type sporangia.

To investigate the role played by PG synthesis, we selected two antibiotics for further characterization: cephalexin, which inhibits PBP activity, and bacitracin, which blocks cell-wall precursor delivery (recycling of undecaprenyl phosphate). Using time-lapse microscopy (see Materials and Methods for details), we monitored membrane dynamics during engulfment in the medial focal plane using the fluorescent membrane dye FM 4-64 (Figure 1B, Video 1). In these 2-5 hour-long movies we observed occasional cell division events occurred with bacitracin (0.08 division events/cell after 90 min, compared to 0.28 division events/cell in untreated cultures, Figure 1-figure supplement 2), indicating that PG synthesis was not completely blocked under these conditions. However, negligible cell divisions occurred with cephalexin, indicating that PG synthesis was indeed completely blocked (Figure 1-figure supplement 2).

To better monitor LE dynamics we used two image analysis approaches (see Materials and Methods for details). First, we created kymographs along forespore membranes (Figure 1E-G). The angular position of forespore pixels was calculated relative to the mother-forespore frame of reference (Figure 1E, inset). All cells were aligned in time based on the onset of septum curving (Figure 1-figure supplement 3), and for a given angle, the average fluorescence of different cells was calculated and plotted over time. Second, we measured the decrease in the distance between the two LEs of the engulfing membrane in the focal plane (the gap arc length), in order to directly assess movement of the LE around the forespore (Figure 1H).

These analyses demonstrated that in untreated sporangia (Figure 1B, top row), the septum curves and the forespore grows into the mother cell without significant forward movement of the LE for ~20 min after polar septation (at 30 °C, Figure 1H). Subsequently, the LE of the engulfing membrane moves towards the forespore pole and engulfment completes within ~60 min (Figure 1E,H). In sporangia treated with cephalexin (Figure 1B, middle row I), the septum curves and extends towards the mother cell, but there is no forward membrane migration (Figure 1F,H). Sometimes the LE retracted on one side while advancing slightly on the other (typically occurred after 90 min of imaging; Figure 1B, middle row II), which appears to be the rotation of the ‘cup’ formed by the engulfing membranes relative to the lateral cell wall (Figure 1C).

Similar to cephalexin, in most sporangia treated with bacitracin (Figure 1B, bottom row I), the forespore extended into the mother cell without significant membrane migration (Figure 1G, H). However, in ~20% of the sporangia, the engulfing membrane migrated asymmetrically, with one side moving faster than the other, although usually it failed to completely surround the forespore (Figure 1B, bottom row II; Figure 1D). The continued engulfment under bacitracin treatment might be related to the fact that PG synthesis is not completely blocked in bacitracin-treated cells under time-lapse conditions (Figure 1-figure supplement 2). Taken together, these results suggest that PG synthesis is not only essential for the final stage of engulfment (membrane fission) in wild type cells [12], but also for migration of the LE of the engulfing membrane around the forespore.

### PBPs accumulate at the leading edge of the engulfing membrane

It has been previously shown that there is an accumulation of membrane-bound PG precursors at the LE of the engulfing membrane [12]. Furthermore, staining with fluorescent d-amino acids has demonstrated that new PG is synthesized at or close to the LE [2]. To investigate if there is a concomitant accumulation of PBPs at the LE, we stained sporangia with BOCILLIN-FL, a commercially available derivative of penicillin V that has a broad affinity for multiple PBPs in *B. subtilis* [28–30]. We observed continuous fluorescent signal around the mother cell membrane that was enriched at the LE (Figure 2A). To better monitor localization of PBPs during engulfment, we plotted fluorescence intensities along the forespores for the membrane and BOCILLIN-FL fluorescent signals as a function of the engulfment stage (Figure 2B). Clearly, the LE is always enriched with PBPs throughout membrane migration.

**Figure 2:**
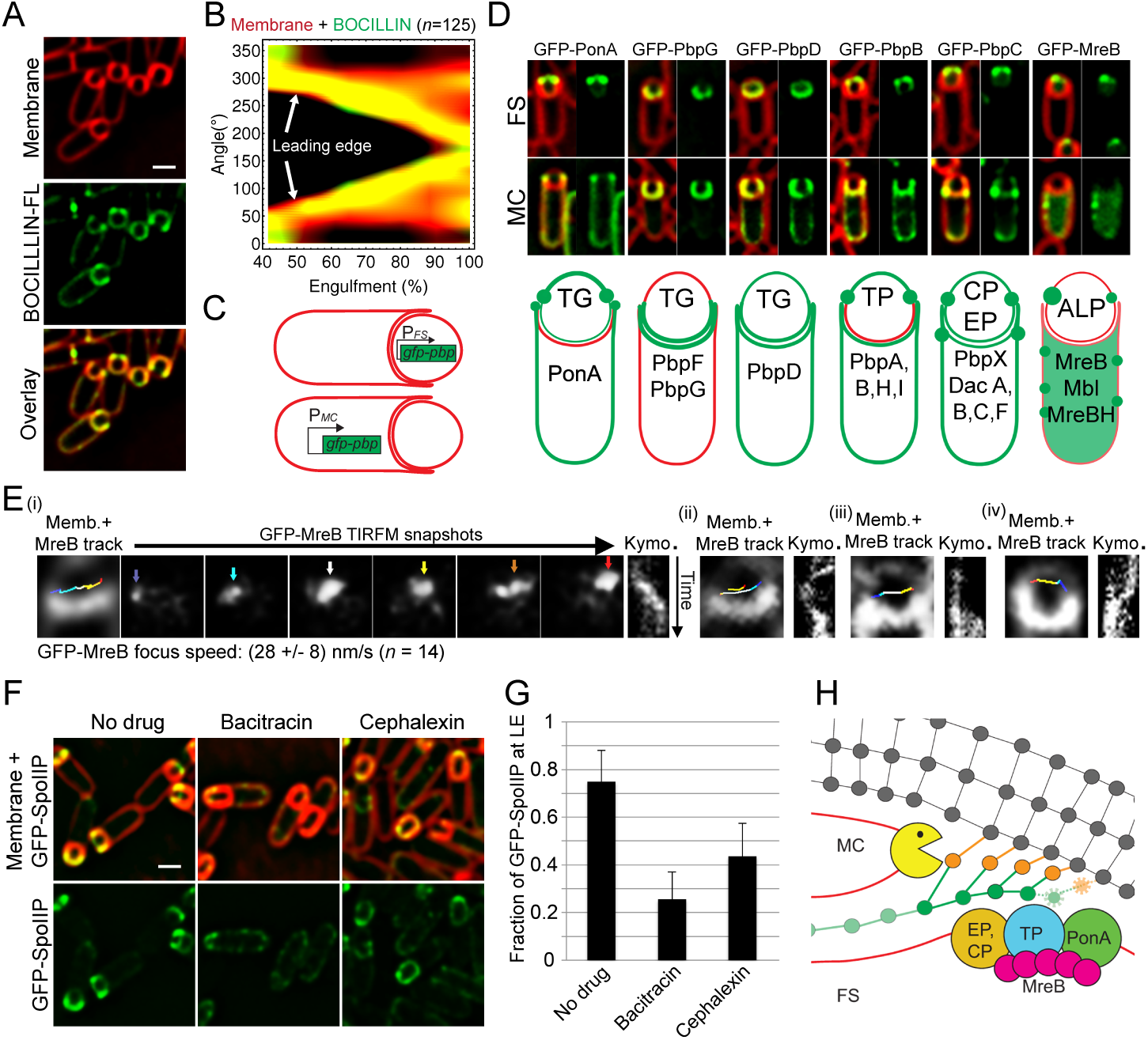
PG synthesis at the LE of the engulfing membrane by forespore PBPs contribute to proper localization of the DMP complex. (A) Sporulating cells stained with a green fluorescent derivative of penicillin V (BOCILLIN-FL). Bright foci are observed at the LE of the engulfing membrane. Membranes were stained with FM 4-64 (red). (B) Average BOCILLIN-FL (green) and FM 4-64 (red) fluorescence intensities along forespore contours plotted as a function of the degree of engulfment. Cells are binned according to percentage of engulfment. BOCILLIN-FL signal is enriched at the LE throughout engulfment (*n* = 125). (C) Cell-specific localization of the peptidoglycan biosynthetic machinery. GFP tagged versions of different *B. subtilis* PBPs and actin-like proteins (ALPs) were produced from mother cell-(MC) or forespore-(FS) specific promoters. (D) Six different localization patterns were observed upon cell-specific localization of PBPs and ALPs. For each pair of images, left panel shows overlay of membrane and GFP fluorescence, while the right panel only shows GFP fluorescence. Pictures of representative cells displaying the different patterns are shown (top, GFP fusion proteins transcribed from *spoIIR* promoter for forespore-specific expression, and from *spoIID* promoter for mother cell-specific expression). The six different patterns are depicted in the bottom cartoon and proteins assigned to each one are indicated. Membranes were stained with FM 4-64. See Figure 2-figure supplement 1 for cropped fields of all PBPs we assayed. Transglycosylase (TG), transpetidase (TP), carboxipetidase (CP), endopeptidase (EP), actin-like protein (ALP). (E) TIRF microscopy of forespore-produced GFP-MreB in four different forespores (i to iv). In every case, the leftmost picture is an overlay of the forespore membranes (shown in white) and the tracks followed by individual TIRF images of GFP-MreB (color encodes time, from blue to red). Sporangia are oriented with the forespores up. For the rst sporangia (i), snapshots from TIRF timelapse experiments taken 8 s apart are shown. Arrows indicate GFP-MreB foci and are color coded to match the trace shown in the left panel. Rightmost panel for each forespore shows a kymograph representing the fluorescence intensity along the line joining the leading edges of the engulfing membrane over time (from top to bottom; total time 100 s). Average focus speed (*n* = 14) is indicated at the bottom. Timelapse movies of the examples presented here and additional sporangia are shown in Video 2. (F) Localizaiton of GFP-SpoIIP in untreated sporangia, or in sporangia treated with bacitracin (50 *μ*g/ml) or cephalexin (50 g/ml). (G) Fraction of GFP-SpoIIP fluorescence at LE of the engulfing membrane. Bars represent the average and standard error of 85 untreated sporangia, 38 sporangia treated with bacitracin (50 *μ*g/ml), and 67 sporangia treated with cephalexin (50 *μ*g/ml). (H) Model for PG synthesis and degradation at the LE of the engulfing membrane. New PG is synthesized ahead of the LE of the engulfing membrane by forespore-associated PG biosynthetic machinery, and is subsequently degraded but the mother-cell DMP complex. We propose that DMP has specificity for the peptide cross-links that join the newly synthesized PG with the lateral cell wall (orange), which leads to the extension of the septal PG around the forespore. Scale bars 1 *μ*m.

### PG biosynthetic machinery tracks the leading edge of the engulfing membrane from the forespore

One possible explanation for the requirement of PG synthesis for engulfment is that PG polymerization by PBPs associated with the LE of the engulfing membrane creates force to pull the engulfing membrane around the forespore. If so, we would expect the PBPs to be located in the mother cell membrane as they polymerize PG. To test this possibility, we assessed the localization of components of the PG biosynthetic machinery in the mother cell or forespore by producing GFP-tagged fusion proteins from promoters that are only active in the mother cell (P_*spoIID*_) or in the forespore (the stronger P_*spoIIQ*_ and the weaker P_*spoIIR*_) after polar septation (Figure 2C,D, Figure 2-figure supplement 1). One prior study tested the localisation of several PBPs during sporulation [31], but most of them were produced before polar separation and it was not possible to determine which cell compartment they were in. We successfully determined the cell-specific localization of 16 proteins involved in PG synthesis (Figure 2-figure supplement 1), including all class A and four class B high-molecular-weight (HMW) PBPs, five low-molecular-weight (LMW) PBPs (four endopeptidases and one carboxipeptidase), and all three MreB paralogues (actin-like proteins, ALPs). Surprisingly, only PonA (PBP1a/b) showed a weak enrichment at the LE of the engul ng membrane when produced in the mother cell (Figure 2D). However, ten PBPs, including PonA and all the class B HMW PBPs and LMW PBPs tested, and all the MreB paralogues were able to track the LE only when produced in the forespore (Figure 2D, Figure 2-figure supplement 1). To follow the dynamics of the forespore PG biosynthetic machinery at the LE, we monitored the movement of GFP-MreB using TIRF microscopy [32, 33]. Forespore GFP-MreB foci rotate around the forespore, coincident with the leading edge of the engulfing membrane, with speeds consistent with those previously reported (Figure 2E, Video 2).

It is unclear how the PBPs recognize the LE, as localization of forespore produced GFP-PonA and GFP-PbpA did not depend on candidate proteins SpoIIB, SpoIID, SpoIIM, SpoIIP, SpoIIQ, SpoIIIAH, SpoIVFAB, or GerM [4, 8, 10, 34, 35] (Figure 2-figure supplement 2). However, these results indicate that the forespore plays a critical role in PG synthesis, and point to an engulfment mechanism that does not depend on pulling the engulfing membrane by mother cell-directed peptidoglycan synthesis.

### PG synthesis is required for SpoIIP localization at the leading edge of the engulfing membrane

The observation that multiple PBPs can track the LE of the engulfing membrane from the forespore opens the possibility that PG synthesis happens ahead of the LE, preceding PG degradation by the mother cell DMP complex. In this context, PG synthesis might be required for proper activity and/or localization of the DMP complex, which is the only other essential engulfment module described so far. The DMP complex localizes at the LE throughout engulfment [9]. To determine if PG synthesis is required for proper localization of DMP, we studied the localization of a GFP-SpoIIP fusion protein when PG synthesis was inhibited by different antibiotics (Figure 2F, G). GFP-SpoIIP shows a well-defined localization at the LE, with ~70% of the total GFP fluorescence at LE in native conditions (Figure 2F, G). However, when PG biosynthesis is inhibited, there is a delocalization of GFP-SpoIIP, which is almost total in cells treated with bacitracin and partial when antibiotics targeting later stages of PG synthesis are used (Figure 2F, G; Figure 2-figure supplement 3). These results are consistent with a model in which PG is synthesized ahead of the LE by forespore-associated PBPs specify the site of PG degradation by the DMP complex (Figure 2G).

### A biophysical model to describe leading edge migration

Our data indicate that engulfment proceeds through coordinated PG synthesis and degradation at the LE. To explain how this coordination leads to engulfment, we propose a minimal biophysical mechanism based on the ‘template mechanism’ of vegetative cell growth assuming that glycans are oriented perpendicular to the long axis of the cell (Figure 3A) [16, 32, 33, 36–38], without requiring any further assumptions about the outer cell wall structure of Gram-positive bacteria, which is still unclear [16, 38, 39]. In this mechanism, a new glycan strand is inserted using both the septal glycan and leading forespore-proximal glycan strand of the lateral wall as template strands to which the new PG strand is cross linked. Subsequently, peptide cross-links between the two template strands are removed from the mother-cell proximal side by the DMP complex. Specifically, in this complex SpoIIP has well documented endopeptidase activity [11]. Note, similar ‘make-before-break’ mechanisms were proposed to allow vegetative cell wall growth without reducing cell wall integrity [36, 37]. A more detailed mechanism that requires the insertion of multiple new glycan strands to account for glycan removal by SpoIID is shown in Figure 3-figure supplement 1. In either model, synthesis of new PG at the LE likely occurs before degradation, thereby naturally preventing cell lysis during engulfment.

**Figure 3:**
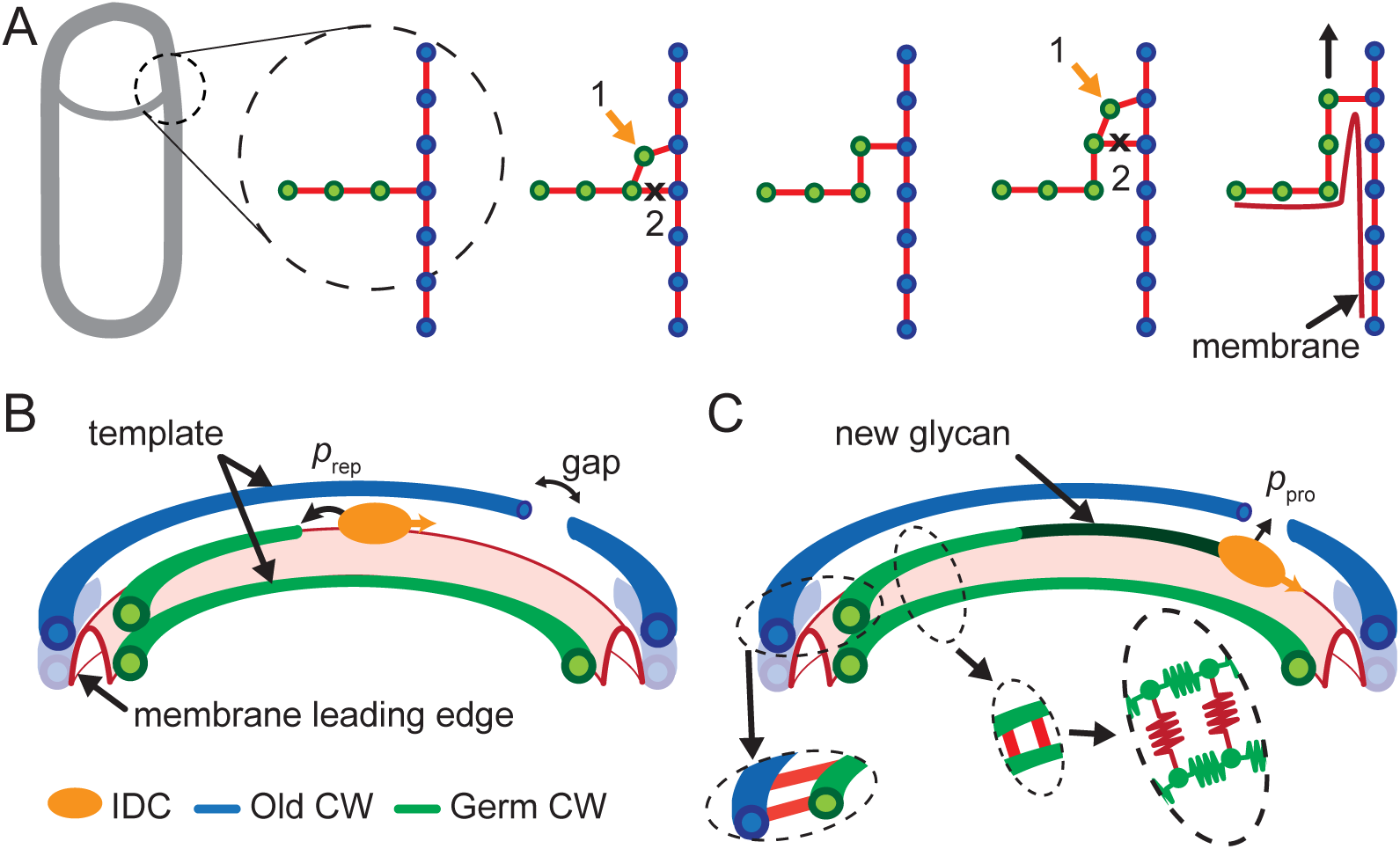
Template model for leading edge (LE) movement. (A) Cell cross-section with glycan strands in the plane perpendicular to the long axis of the cell. One strand from old cell wall (blue) and one strand from newly synthesized germ-cell wall (green) are used as a template for new glycan insertion. Coordination between glycan insertion (orange arrow) and peptide cross-link degradation (black cross) drives LE forward. (B) 3D model of stochastic glycan insertion by insertion-degradation complex (IDC) with transpeptidase and transglycosylase activity. Probability of IDC to start inserting new glycan from old glycan end and repair end defect is *p*_rep_. (C) New inserted glycan shown in dark green. Probability of IDC to continue glycan insertion when it encounters gap in old cell wall is probability of processivity *p*_pro_. (Inset) Horizontal (between old and new glycan strands) and vertical (between new glycan strands) peptide links are shown in red. In our coarse-grained model glycans are simulated as semi-flexible filaments consisting of beads (green) connected with springs (green). Peptides are simulated as springs (red) connecting neighboring glycan beads.

The coordination between PG insertion from the forespore and removal by DMP in the mother cell could lead to movement of the junction between the septal peptidoglycan and the lateral peptidoglycan around the forespore to mediate successful engulfment. Based on this proposed mechanism, we created a model whereby insertion and degradation happens, for simplicity, simultaneously by an insertion-degradation complex (IDC), also reflecting the high degree of coordination suggested by the template mechanism. In this model IDC recognizes the leading edge and inserts glycan polymers perpendicular to the long axis of the cell (Figure 3B). Additionally, the model proposes that IDC can recognize glycan ends and initiate glycan polymerization from the end defect with probability of repair *p*_rep_. During glycan insertion, when an IDC encounters a gap in the outer cell wall strands, it continues polymerization with probability of processivity *p*_pro_ (Figure 3C). A systematic exploration of the above model parameters showed that intact spores form for *p*_rep_ and *p*_pro_ > 0.8 with a marginal dependence on the number of IDCs (Figure 4G, Figure 4-figure supplement 1). However, to compare the model with microscopy data we require a 3D dynamic implementation of this model that reflects the stochasticity of underlying molecular events.

**Figure 4:**
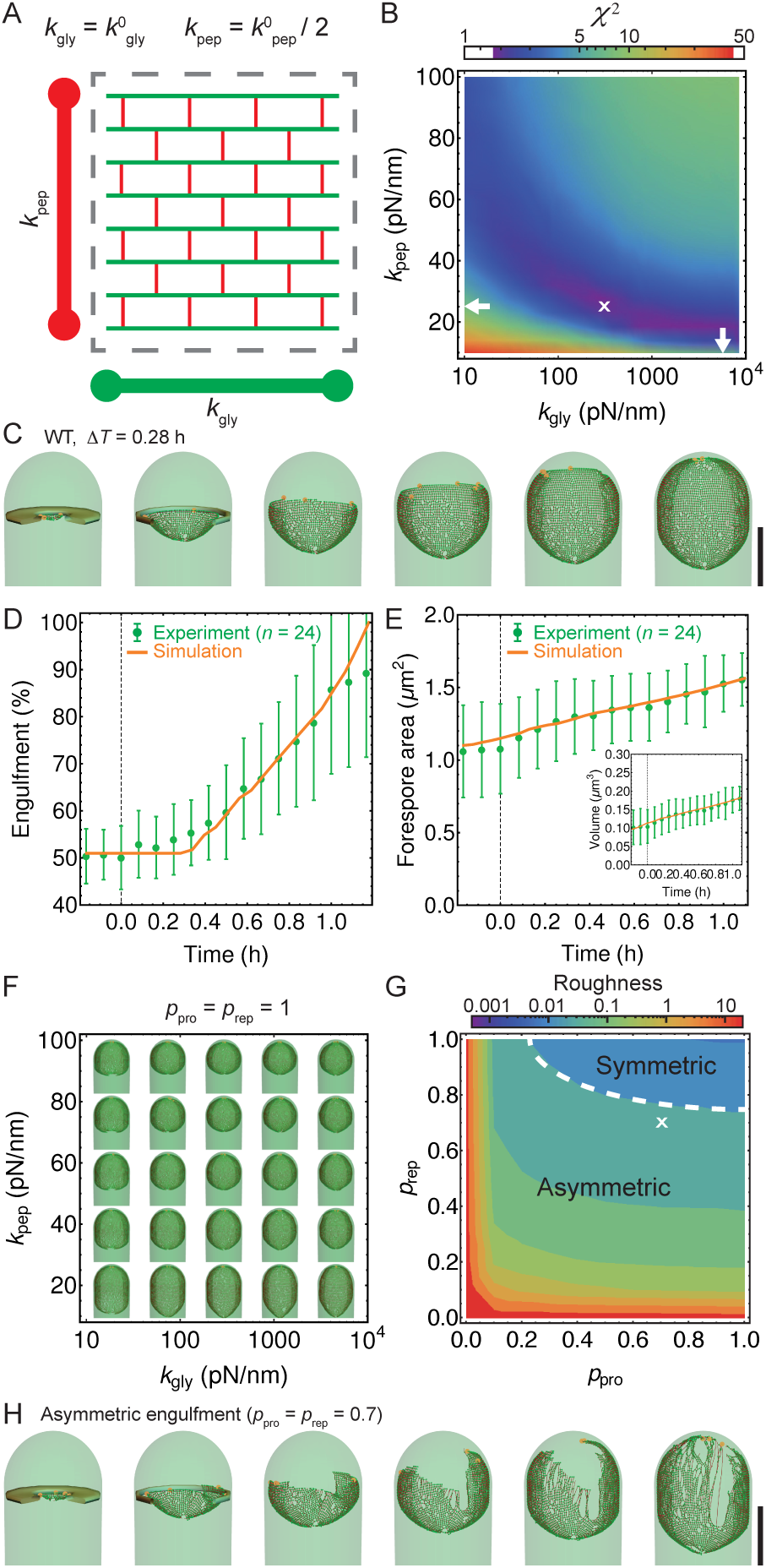
Template model reproduces experimentally observed phenotypes. (A) Effective spring constants in our model represent coarse-grained PG network. Here the angle between neighboring stem peptides that belong to a single glycan is assumed to be 90°. Therefore, every other stem peptide is in plane with glycan sheet [42, 43]. The role of effective glycan persistence length on engulfment is negligible (see Figure 4-figure supplement 3). (B) Simulations for different values of effective peptide *k*_pep_ and glycan *k*_gly_ spring constants are compared with experimentally measured forespore surface area, volume and engulfment using mutual *χ*^2^ statistics (Eq. 2). Arrows point to effective literature *k*_pep_ and *k*_gly_ [43]. Dark blue region corresponds to simulation parameters that best fit experimental data (Figure 4-figure supplement 4, Video 3). For large enough *k*_gly_ > 200 pN/nm mutual *χ*^2^ is almost independent of *k*_gly_. (C) Snapshots of WT simulations for parameters (*k*_gly_ = 200 pN/nm, *k*_pep_ = 25 pN/nm, *N*_IDC_ = 5) marked with ‘×’ in panel (B) (Video 2). The thick septum is treated as outer cell wall, and is assumed degraded once IDCs move along. (D-E) Time traces of experimentally measured engulfment, forespore surface area and forespore volume (green) in comparison with results from a single simulation (orange). Parameters used in simulation are marked with ‘×’ in panel (B). For all other parameters see *SI Appendix*. (F) Snapshots of fully engulfed forespores for various peptidoglycan elastic constants. (G) For various values of independent parameters *p*_rep_ and *p*_pro_ roughness of the LE is calculated at the end of stochastic simulations (see Figure 4-figure supplement 1, and Video 4). Here 0 roughness correspond to perfectly symmetric LE; for high enough *p*_rep_ = *p*_pro_ > 0.8 LE forms symmetric pro les. (H) Simulation for asymmetric engulfment is obtained for same parameter as WT except *p*_rep_ = *p*_pro_ = 0.7 (marked with ‘×’ in panel (G)). Average ± SD. Scale bars 1 *μ*m.

### Langevin simulations reproduce observed phenotypes

To simulate stochastic insertion at the leading edge we used Langevin dynamics of a coarse-grained PG meshwork (see Materials and Methods). Briefly, glycan strands are modeled as semi-flexible filaments consisting of beads connected with springs, while peptide bridges are modeled as springs connecting glycan beads (Figure 3C) [40–42]. Typical length of inserted glycan polymer is ~1 *μ*m (~1/3 cell circumference) [39] and in our model the peptide bridges between newly inserted glycan strands are in a relaxed state. Glycan beads experience forces due to glycan elastic springs (*k*_gly_), glycan persistence length (*l*_p_), elastic peptide links (*k*_pep_), stochastic thermal fluctuations, and pressure difference (Δ*p*) between forespore and mother cell (see Eq. 1 and *SI Appendix*). Glycan strands in the PG layer are connected with neighboring glycans by stem peptides (Figure 4A). In our model, the angle between neighboring stem peptides that belong to the same glycan strand is assumed to be 90° [42, 43]. Therefore, every other stem peptide is in plane with the glycan sheet. In our model Δ*p* originates from the packing of the *B. subtilis* chromosome (~4.2 Mbp) in the small forespore compartment [44–47].

To systematically explore the peptidoglycan parameters, we compared our simulations with actual changes in forespore volume, forespore surface area, and percentage of engulfment extracted from time-lapse movies, using χ^2^ fitting (Figure 4B, Eq. 2, Materials and Methods). Parameters that best fit experimental measurements belong to dark blue region in agreement with molecular dynamic simulations [43]. For a single peptide bond, the linear elasticity regime is valid for extensions that are less than 1 nm [43] and this elastic regime is maintained in the regions with low χ^2^ (Figure 4-figure supplement 2). For large enough glycan stiffness (*k*_gly_ > 300 pN/nm)*χ*^2^ becomes independent of *k*_gly_ (Figure 4B). A typical simulation shown in Figure 4C matches experimental measurements of time-dependent engulfment, volume, and surface area (Figure 4D, E). PG spring constants drastically affect forespore morphologies. By decreasing *k*_pep_ forespores elongate, while by increasing *k*_pep_ forespores shrink, as measured along the long axis of the cell. Changing *k*_gly_ has only minor effects on volume and surface area. However, the main effect is on forespore curvature (see Figure 4-figure supplement 4): high *k*_gly_ increases the curvature of forespore ends (making them more pointy), while low *k*_gly_ decreases the curvature of the forespore ends. Our simulations successfully reproduce asymmetric engulfment (Figure 4F, G; Video 5). For *p*_rep_ and *p*_pro_ ⩽ 0.8 we obtained asymmetric engulfment that reproduces the phenotypes observed when PG synthesis or degradation is partially blocked. When defects in the peptidoglycan meshwork are not repaired, different parts of the leading edge extend in an uncoordinated manner, producing asymmetric engulfment.

Since our simulations correctly reproduced engulfment dynamics we used simulation parameters to estimate glycan insertion velocities *V*_IDC_ of IDC (see *SI Appendix*). Using this method we estimated a lower bound on product *N*_IDC_ · *V*_IDC_ ~110 nm/s, where *N*_IDC_ is the number of insertion complexes. Similarly, by estimating the total amount of newly inserted material in the forespore within ~0.8 h without any pausing we obtain *N*_IDC_ · *V*_IDC_ ~117 nm/s. For circumferentially processive PBPs (PbpA and PbpH), the absolute velocity measured using TIRF microscopy is ~20-40 nm/s during vegetative cell growth [32, 33], which is in agreement with the speed of forespore GFP-MreB determined from our TIRF experiments ((28 ± 8) nm/s, *n* = 14; Figure 2E). Using this estimate for *V*_IDC_, we obtain a lower bound 3-6 on the number of active, highly processive PBP molecules. However, the actual number of proteins could be higher for other nonprocessive PBPs [32, 33].

## Discussion

The results presented here suggest that engulfment involves coordinated PG synthesis and degradation processes that are segregated between different cell types: first, PG is synthesized in front of the LE of the engulfing membrane by a forespore-associated PG biosynthetic machinery that rotates following the LE of the engulfing membrane. Then this new PG is targeted for degradation by the mother cell-associated PG degradation machinery comprised of the DMP complex (Figure 2H). The delocalization of DMP when PG synthesis is inhibited with antibiotics (Figure 2, Figure 2-figure supplement 3) indicates that the DMP either forms an actual complex with the PG biosynthetic machinery across the septal PG (to form a single insertion degradation complex (IDC), as shown in Fig. 3) or that DMP targets the new PG synthesized at the LE of the engulfing membrane. In the latter, DMP might specifically target the cross-links that attach the old lateral cell wall to the new PG synthesized at the LE of the engulfing membrane (Figure 2H, orange). Since those cross-links join old, modified PG from the lateral cell wall to newly synthesized PG at the LE, those peptide bridges might have a unique chemical composition or structural arrangement that could be specifically recognized by DMP. Hence, either approach provides a safety mechanism during engulfment, since it would prevent DMP from degrading the old PG of the lateral cell wall, which could lead to cell lysis.

We have conceptualized these results in a biophysical model in which a PG insertion-degradation complex (IDC), representing PBPs for PG synthesis and DMP proteins for PG degradation, catalyzes PG remodeling at the LE of the engulfing membrane. Specifically, we propose that new glycan strands are inserted ahead of the LE of the engulfing membrane and PG is degraded on the mother cell proximal side to create space for forward movement of the LE (Figure 3). This is similar to the ‘make-before-break’ model of vegetative cell-wall growth, which postulates that the vegetative cell wall is elongated by inserting new PG strands prior to degrading old strands [36] (although bacteria can also make a de novo cell wall [48, 49]). The make-before-break mechanism also accounts for the directional movement of the LE towards the forespore pole, since the substrate for DMP is new PG synthesized by forespore PBPs, which is always ahead of the LE of the engulfing membrane.

Using Langevin simulations we successfully reproduced the dynamics of engulfment, forespore volume, and surface area. Additionally, our model correctly reproduced asymmetric engulfment observed with reduced IDC activity, and we estimated that with only a handful of highly processive PBP molecules are necessary to reproduce the observed LE dynamics. A more general model without strong coupling between the PG biosynthetic and PG degradation machineries also leads to successful engulfment (SI Appendix, Figure 4-figure supplement 5, Video 6). However, DMP has to be guided to degrade only the peptide cross-links between old and new glycan strands, and should also prevent detachment of the septal peptidoglycan from the old cell wall.

Since our simple mechanism in Figure 3A entails hydrolysis of certain peptide bonds but no glycan degradation, we explored additional mechanisms since the SpoIID protein of the DMP complex shows transglycosylase activity [11]. First, it is possible that engulfment entails a two-for-one mechanism, with two new glycan strands are added and the newly inserted glycan strand at the LE is degraded [37] (Figure 3-figure supplement 1A). Similarly, the three-for-one mechanism would also work [50]. Second, one new glycan strand might be added and the inner most cell-wall glycan of the thick, lateral cell wall degraded (Figure 3-figure supplement 1B). This would make the lateral cell wall thinner as the engulfing membrane moves forward [2]. Finally, it is possible that insertion and degradation are not intimately coupled, and that there is simply a broad region in which PG is inserted ahead of the engulfing membrane, to create multiple links between the septal PG and the lateral cell wall (as shown in Figure 2H), and that the DMP complex has a preference for newly synthesized PG. All of these models require the spatial coordination between cell wall degradation and synthesis to avoid compromising cell wall integrity and inducing cell lysis, and all share a common ‘make-before-break’ strategy to promote robustness of the otherwise risky PG remodeling process [36]. In order to waste as little energy as possible, a more stringent ‘make-just-before-break’ strategy may even apply, motivating more intimate coupling between the PG biosynthetic and degradation machineries.

Our simple biophysical mechanism postulates that engulfment does not rely on pulling or pushing forces for membrane migration. Instead, cell wall remodeling makes room for the mother cell membrane to expand around the forespore by entropic forces. During engulfment the mother-cell surface area increases by ~2 *μ*m^2^ (~25 %, see Figure 1-figure supplement 3), and this excess of membrane could simply be accommodated around the forespore by remodeling the PG at the LE. However, our model does not include all potential contributors to engulfment. For instance, the SpoIIQ-AH zipper, which is dispensable for engulfment in native conditions [5], might prevent membrane backward movement, and might also help localize the IDC components toward the LE. Interestingly, SpoIIQ-AH interaction is essential for engulfment in *Clostridium difficile* where the SpoIIQ ortholog posseses endopeptidase activity [51–53]. The model also does not consider the impact of the tethering of the LE of the engulfing membrane to the forespore via interactions between the mother cell membrane anchored DMP complex at the LE and forespore synthesized PG. Future experiments and modeling should address the role of these and other potential contributors to LE migration, which will allow us to re ne our biophysical model and obtain a comprehensive view of membrane dynamics during engulfment. Furthermore, understanding the cooperation between PBPs and DMP will provide valuable clues about the structure of the cell wall in Gram-positive bacteria.

## Materials and Methods

### Strains and culture conditions

All the strains used in this study are derivatives of *B. subtilis* PY79. Complete lists of strains and plasmids, as well as detailed descriptions of plasmid construction are provided in *SI Appendix*. For each experiment we had at least two biological replicas, and each one contains at least three technical replicas. Averages of individual cells, but not the averages of different replicas are reported. Sporulation was induced by resuspension [54], except that the bacteria were grown in 25 % LB prior to resuspension, rather than CH medium. Cultures were grown at 37 °C for batch culture experiments, and at 30 °C for timelapse experiments.

### Fluorescence microscopy

Cells were visualized on an Applied Precision DV Elite optical sectioning microscope equipped with a Photometrics CoolSNAP-HQ^2^ camera and deconvolved using SoftWoRx v5.5.1 (Applied Precision). When appropriate, membranes were stained with 0.5 *μ*g/ml FM 4-64 (Life Technologies) or 1 *μ*g/ml Mitotracker green (Life Technologies). Cells were transferred to 1.2 % agarose pads for imaging. The median focal plane is shown.

### Deconvolution fluorescent microscopy

Sporulation was induced at 30 °C. 1.5 h after sporulation induction, 0.5 *μ*g/ml FM 4-64 was added to the culture and incubation continued for another 1.5 h. Seven *μ*l samples were taken 3 h after resuspension and transferred to agarose pads prepared as follows: 2/3 volume of supernatant from the sporulation culture; 1/3 volume 3.6 % agarose in fresh A+B sporulation medium; 0.17 *μ*g/ml FM 4-64. When appropriated, cephalexin (50 *μ*g/ml) or bacitracin (50 *μ*g/ml) was added to the pad. Pads were partially dried, covered with a glass slide and sealed with petroleum jelly to avoid dehydration during timelapse imaging. Petroleum jelly is not toxic and cannot be metabolized by *B. subtilis*, which poses an advantage over other commonly used sealing compounds, such as glycerol, which can be used as a carbon source and inhibit the initiation of sporulation. Pictures were taken in an environmental chamber at 30 °C every 5 min for 5 h. Excitation/emission filters were TRITC/CY5. Excitation light transmission was set to 5 % to minimize phototoxicity. Exposure time was 0.1 s.

### Forespore GFP-MreB tracking experiments

MreB tracking experiments were performed using the strain JLG2411, which produced GFP-MreB in the forespore after polar septation from *spoIIQ* promoter. Sporulation and agarose pads were done as described in *Deconvolution fluorescent microscopy*, except that FM 4-64 was only added to the agarose pads and not to the sporulating cultures. A static membrane picture was taken at the beginning of the experiment, and was used as a reference to determine the position of the GFP-MreB foci. GFP-MreB motion at the cell surface was determined by TIRF microscopy [32, 33], taking pictures every 4 s for 100 s. Imaging was performed at 30 °C. We used two different microscopes to perform TIRF microscopy: (i) An Applied Precision Spectris optical sectioning microscope system equipped with an Olympus IX70 microscope, a Photometrics CoolSNAP HQ digital camera and a 488-nm argon laser. To perform TIRF in this microscope, we used an Olympus 1003 1.65 Apo objective, immersion oil *n* = 1.78 (Cargille Laboratories), and sapphire coverslips (Olympus). Laser power was set to 15 %, and exposure time was 200 ms. (ii) An Applied Precision OMX Structured Illumination microscopy equipped with a Ring-TIRF system and a UApoN 1.49NA objective, immersion oil *n* = 1.518. Exposure time was 150 ms.

Images were analyzed using the ImageJ-based FIJI package. Sporangia were aligned vertically using the rotation function in FIJI. GFP-MreB foci were tracked using the TrackMate pluging [55], using the LoG detector, estimated blob diameter of 300 nm, simple LAP tracked and linking max distance of 300 nm. Only tracks that contained more than four points were used to determine the MreB foci speed.

### Image analysis

We used the semi-automated active contour software JFilament available as ImageJ plugin to extract fluorescently labeled membrane position over time [56]. Membrane position obtained from the medial focal plane is used in custom built *Mathematica* software to calculate 3D volume and surface area by assuming rotational symmetry around the axis connecting the center of masses of mother cell and forespore. For available code and example see supplementary file ‘image analysis example:zip’. Kymographs as in Figure 1E were created by collecting intensities along the forespore contours. Subsequently, pixel angles were determined using pixel position relative to the mother-forespore frame as defined in inset of Figure 1E. Forespore fluorescent intensities along angles are normalized and interpolated using third-order polynomials. For a given angle the population intensity average of different cells is calculated and plotted over time. Time 0’ is the onset of septum curving.

### Quanti cation of GFP-SpoIID, GFP-SpoIIM and GFP-SpoIIP fraction at LE

Antibiotics were added 2 hours after resuspension, and samples were taken one hour later for imaging. Exposure times and image adjustments were kept constant throughout the experiment. To determine the fraction of GFP signal at the LE, GFP pixel intensities of seven optical sections covering a total thickness of 0.9 *μ*m were summed. GFP intensities at the LE (*I*_LE_) and in the rest of the mother cell (*I*_MC_) were determined separately by drawing polygons encompassing the LE or the MC. After subtraction of the average background intensity, the fraction of GFP fluorescence at 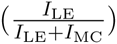 was determined for each sporangium.

### Langevin dynamics

The Langevin dynamic equation of the *i*^th^ bead at position **r**_*i*_ is given by:

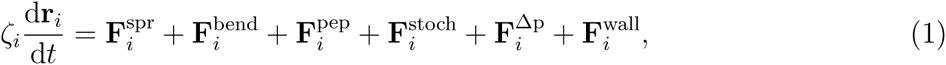

where the left-hand side depends on the drag coefficient *ζ_i_* ≈ 4π*η*_med_*l*_0_ [57], with *η*_med_ is the medium viscosity and *l*_0_ equilibrium distance between neighbouring beads (see *SI Appendix*). On the right-hand side of Eq. 1 we have contributions of glycan elastic spring, glycan bending, peptide elastic links, stochastic thermal fluctuations, pressure difference Δ*p* between forespore and mother, and excluded volume from the old cell wall, respectively.

### *χ*^2^ tting of parameters

To compare simulations with experiments we measured forespore volume (*V_i_*), forespore surface area (*S_i_*) and engulfment (*E_i_*) and constructed a quality-of-fit function as:

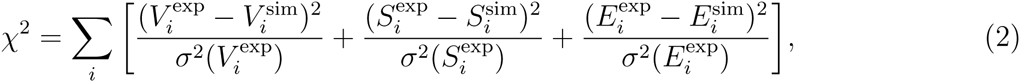

where index *i* corresponds to *i*^th^ time point, and σ is the standard deviation [58].

Details are given in the *SI Appendix*.

## Acknowledgments

This work is supported by European Research Council Starting Grant 280492-PPHPI to NO and RGE, EMBO Long Term Fellowship to JLG, National Institutes of Health R01-GM57045 to KP. We thank the Microscopy Core at UC San Diego (P30 NS047101) for help in using the Applied Precision OMX Structured Illumination microscopy used for TIRF experiments. The funders had no role in study design, data collection and analysis, decision to publish, or preparation of the manuscript.

## References

[1] Tan, I. S & Ramamurthi, K. S. (2014) Spore formation in Bacillus subtilis. Environ Microbiol Rep 6, 212–225.

[2] Tocheva, E. I, López-Garrido, J, Hughes, H, Fredlund, J, Kuru, E, VanNieuwenhze, M. S, Brun, Y. V, Pogliano, K, & Jensen, G. J. (2013) Peptidoglycan transformations during Bacillus subtilis sporulation. Mol Microbiol 88, 673–686.

[3] Higgins, D & Dworkin, J. (2012) Recent progress in Bacillus subtilis sporulation. FEMS Microbiol Rev 36, 131–148.

[4] Blaylock, B, Jiang, X, Rubio, A, Moran, C. P, & Pogliano, K. (2004) Zipper-like interaction between proteins in adjacent daughter cells mediates protein localization. Genes Dev 18, 2916–2928.

[5] Broder, D. H & Pogliano, K. (2006) Forespore engulfment mediated by a ratchet-like mechanism. Cell 126, 917–928.

[6] Ojkic, N, López-Garrido, J, Pogliano, K, & Endres, R. G. (2014) Bistable forespore engulfment in Bacillus subtilis by a zipper mechanism in absence of the cell wall. PLoS Comput Biol 10, e1003912.

[7] Sun, Y.-L, Sharp, M. D, & Pogliano, K. (2000) A dispensable role for forespore-specific gene expression in engulfment of the forespore during sporulation ofbacillus subtilis. J Bacteriol 182, 2919–2927.

[8] Abanes-De Mello, A, Sun, Y.-L, Aung, S, & Pogliano, K. (2002) A cytoskeleton-like role for the bacterial cell wall during engulfment of the Bacillus subtilis forespore. Genes Dev 16, 3253–3264.

[9] Gutierrez, J, Smith, R, & Pogliano, K. (2010) SpoIID-mediated peptidoglycan degradation is required throughout engulfment during Bacillus subtilis sporulation. J Bacteriol 192, 3174–3186.

[10] Chastanet, A & Losick, R. (2007) Engulfment during sporulation in Bacillus subtilis is governed by a multi-protein complex containing tandemly acting autolysins. Mol Microbiol 64, 139–152.

[11] Morlot, C, Uehara, T, Marquis, K. A, Bernhardt, T. G, & Rudner, D. Z. (2010) A highly coordinated cell wall degradation machine governs spore morphogenesis in Bacillus subtilis. Genes Dev 24, 411–422.

[12] Meyer, P, Gutierrez, J, Pogliano, K, & Dworkin, J. (2010) Cell wall synthesis is necessary for membrane dynamics during sporulation of Bacillus subtilis. Mol Microbiol 76, 956–970.

[13] Reith, J & Mayer, C. (2011) Peptidoglycan turnover and recycling in Gram-positive bacteria. Appl Microbiol Biot 92, 1–11.

[14] Lee, T. K & Huang, K. C. (2013) The role of hydrolases in bacterial cell-wall growth. Curr Opin Microbiol 16, 760–766.

[15] Misra, G, Rojas, E. R, Gopinathan, A, & Huang, K. C. (2013) Mechanical consequences of cell-wall turnover in the elongation of a Gram-positive bacterium. Biophys J 104, 2342–2352.

[16] Dover, R. S, Bitler, A, Shimoni, E, Trieu-Cuot, P, & Shai, Y. (2015) Multiparametric AFM reveals turgor-responsive net-like peptidoglycan architecture in live streptococci. Nat Commun 6, 7193.

[17] Tocheva, E. I, Matson, E. G, Morris, D. M, Moussavi, F, Leadbetter, J. R, & Jensen, G. J. (2011) Peptidoglycan remodeling and conversion of an inner membrane into an outer membrane during sporulation. Cell 146, 799–812.

[18] Grant, W. (1979) Cell wall teichoic acid as a reserve phosphate source in Bacillus subtilis. J Bacteriol 137, 35–43.

[19] Brown, S, Santa Maria Jr, J. P, & Walker, S. (2013) Wall teichoic acids of gram-positive bacteria. Annu Rev Microbiol 67.

[20] Chin, T, Younger, J, & Glaser, L. (1968) Synthesis of teichoic acids during spore germination. J Bacteriol 95, 2044–2050.

[21] Johnstone, K, Simion, F. A, & Ellar, D. J. (1982) Teichoic acid and lipid metabolism during sporulation of Bacillus megaterium KM. Biochem J 202, 459–467.

[22] Lizunov, V & Zimmerberg, J. (2006) Cellular biophysics: Bacterial endospore, membranes and random fluctuation. Curr Biol 16, R1025–R1028.

[23] Pogliano, J, Osborne, N, Sharp, M. D, Mello, A.-D, Perez, A, Sun, Y.-L, Pogliano, K, et al. (1999) A vital stain for studying membrane dynamics in bacteria: a novel mechanism controlling septation during Bacillus subtilis sporulation. Mol Microbiol 31, 1149–1159.

[24] González-Pastor, J. E, Hobbs, E. C, & Losick, R. (2003) Cannibalism by sporulating bacteria. Science 301, 510–513.

[25] Straight, P. D & Kolter, R. (2009) Interspecies chemical communication in bacterial development. Annu Rev Microbiol 63, 99–118.

[26] Lamsa, A, Liu, W.-T, Dorrestein, P. C, & Pogliano, K. (2012) The Bacillus subtilis cannibalism toxin SDP collapses the proton motive force and induces autolysis. Mol Bicrobiol 84, 486–500.

[27] Lamsa, A, Lopez-Garrido, J, Quach, D, Riley, E. P, Pogliano, J, & Pogliano, K. (2016) Rapid inhibition profiling in Bacillus subtilis to identify the mechanism of action of new antimicrobials. ACS Chem Biol.

[28] Lakaye, B, Damblon, C, Jamin, M, Galleni, M, Lepage, S, Joris, B, Marchand-Brynaert, J, Frydrych, C, & Frere, J.-M. (1994) Synthesis, purification and kinetic properties of fluorescein-labelled penicillins. Biochem J 300, 141–145.

[29] Zhao, G, Meier, T. I, Kahl, S. D, Gee, K. R, & Blaszczak, L. C. (1999) BOCILLIN FL, a sensitive and commercially available reagent for detection of penicillin-binding proteins. Antimicrob Agents Ch 43, 1124–1128.

[30] Kocaoglu, O, Calvo, R. A, Sham, L.-T, Cozy, L. M, Lanning, B. R, Francis, S, Winkler, M. E, Kearns, D. B, & Carlson, E. E. (2012) Selective penicillin-binding protein imaging probes reveal substructure in bacterial cell division. ACS Chem Biol 7, 1746–1753.

[31] Scheffers, D.-J. (2005) Dynamic localization of penicillin-binding proteins during spore development in Bacillus subtilis. Microbiol 151, 999–1012.

[32] Garner, E. C, Bernard, R, Wang, W, Zhuang, X, Rudner, D. Z, & Mitchison, T. (2011) Coupled, circumferential motions of the cell wall synthesis machinery and MreB filaments in B. subtilis. Science 333, 222–225.

[33] Domínguez-Escobar, J, Chastanet, A, Crevenna, A. H, Fromion, V, Wedlich-Söldner, R, & Carballido-López, R. (2011) Processive movement of mreb-associated cell wall biosynthetic complexes in bacteria. Science 333, 225–228.

[34] Aung, S, Shum, J, Mello, A.-D, Broder, D. H, Fredlund-Gutierrez, J, Chiba, S, & Pogliano, K. (2007) Dual localization pathways for the engulfment proteins during Bacillus subtilis sporulation. Mol Microbiol 65, 1534–1546.

[35] Rodrigues, C. D, Ramírez-Guadiana, F. H, Meeske, A. J, Wang, X, & Rudner, D. Z. (2016) GerM is required to assemble the basal platform of the SpoIIIA–SpoIIQ transenvelope complex during sporulation in Bacillus subtilis. Mol Microbiol.

[36] Koch, A. L & Doyle, R. J. (1985) Inside-to-outside growth and turnover of the wall of gram-positive rods. J Theor Biol 117, 137–157.

[37] Höltje, J.-V. (1998) Growth of the stress-bearing and shape-maintaining murein sacculus of Escherichia coli. Microbiol Mol Biol Rev 62, 181–203.

[38] Beeby, M, Gumbart, J. C, Roux, B, & Jensen, G. J. (2013) Architecture and assembly of the Gram-positive cell wall. Mol Microbiol 88, 664–672.

[39] Hayhurst, E. J, Kailas, L, Hobbs, J. K, & Foster, S. J. (2008) Cell wall peptidoglycan architecture in Bacillus subtilis. Proc Natl Acad Sci USA 105, 14603–14608.

[40] Laporte, D, Ojkic, N, Vavylonis, D, & Wu, J.-Q. (2012) *α*-actinin and fimbrin cooperate with myosin II to organize actomyosin bundles during contractile-ring assembly. Mol Biol Cell 23, 3094–3110.

[41] Tang, H, Laporte, D, & Vavylonis, D. (2014) Actin cable distribution and dynamics arising from cross-linking, motor pulling, and filament turnover. Mol Biol Cell 25, 3006–3016.

[42] Huang, K. C, Mukhopadhyay, R, Wen, B, Gitai, Z, & Wingreen, N. S. (2008) Cell shape and cell-wall organization in Gram-negative bacteria. Proc Natl Acad Sci USA 105, 19282–19287.

[43] Nguyen, L. T, Gumbart, J. C, Beeby, M, & Jensen, G. J. (2015) Coarse-grained simulations of bacterial cell wall growth reveal that local coordination alone can be sufficient to maintain rod shape. Proc Natl Acad Sci USA 112, E3689–E3698.

[44] Errington, J. (1993) Bacillus subtilis sporulation: regulation of gene expression and control of morphogenesis. Microbiol Rev 57, 1–33.

[45] Perez, A. R, Abanes-De Mello, A, & Pogliano, K. (2000) SpoIIB localizes to active sites of septal biogenesis and spatially regulates septal thinning during engulfment in Bacillus subtilis. J Bacteriol 182, 1096–1108.

[46] Bath, J, Wu, L. J, Errington, J, & Wang, J. C. (2000) Role of Bacillus subtilis SpoIIIE in DNA transport across the mother cell-prespore division septum. Science 290, 995–997.

[47] Shin, J. Y, Lopez-Garrido, J, Lee, S.-H, Diaz-Celis, C, Fleming, T, Bustamante, C, & Pogliano, K. (2015) Visualization and functional dissection of coaxial paired SpoIIIE channels across the sporulation septum. eLife 4, e06474.

[48] Ranjit, D. K & Young, K. D. (2013) The Rcs stress response and accessory envelope proteins are required for de novo generation of cell shape in Escherichia coli. J Bacteriol 195, 2452–2462.

[49] Kawai, Y, Mercier, R, & Errington, J. (2014) Bacterial cell morphogenesis does not require a preexisting template structure. Curr Biol 24, 863–867.

[50] Scheffers, D.-J & Pinho, M. G. (2005) Bacterial cell wall synthesis: new insights from localization studies. Microbiol Mol Biol Rev 69, 585–607.

[51] Crawshaw, A. D, Serrano, M, Stanley, W. A, Henriques, A. O, & Salgado, P. S. (2014) A mother cell-to-forespore channel: current understanding and future challenges. FEMS Microbiol Lett 358, 129–136.

[52] Serrano, M, Crawshaw, A. D, Dembek, M, Monteiro, J. M, Pereira, F. C, de Pinho, M. G, Fairweather, N. F, Salgado, P. S, & Henriques, A. O. (2015) The SpoIIQ-SpoIIIAH complex of Clostridium difficile controls forespore engulfment and late stages of gene expression and spore morphogenesis. Mol Microbiol 100, 204–228.

[53] Fimlaid, K. A, Jensen, O, Donnelly, M. L, Siegrist, M. S, & Shen, A. (2015) Regulation of Clostridium difficile spore sormation by the SpoIIQ and SpoIIIA proteins. PLoS Genet 11, e1005562.

[54] Sterlini, J. M & Mandelstam, J. (1969) Commitment to sporulation in Bacillus subtilis and its relationship to development of actinomycin resistance. Biochem J 113, 29–37.

[55] Tinevez, J.-Y, Perry, N, Schindelin, J, Hoopes, G. M, Reynolds, G. D, Laplantine, E, Bednarek, S. Y, Shorte, S. L, & Eliceiri, K. W. (2016) TrackMate: An open and extensible platform for single-particle tracking. Methods.

[56] Smith, M. B, Li, H, Shen, T, Huang, X, Yusuf, E, & Vavylonis, D. (2010) Segmentation and tracking of cytoskeletal filaments using open active contours. Cytoskeleton 67, 693–705.

[57] Howard, J. (2001) Mechanics of motor proteins and the cytoskeleton. (Sinauer Associates).

[58] Spitzer, P, Zierhofer, C, & Hochmair, E. (2006) Algorithm for multi-curve-fitting with shared parameters and a possible application in evoked compound action potential measurements. Biomed Eng Online 5, 13.

